# Cell morphology and gene expression: tracking changes and complementarity across time and cell lines

**DOI:** 10.1101/2024.08.30.610494

**Authors:** Vanille Lejal, David Rouquié, Olivier Taboureau

## Abstract

Effective drug discovery relies on combining target knowledge with functional assays and multi-omics data to understand chemicals’ molecular actions. However, the relationship between cell morphology and gene expression over time and across cell lines remains unclear. To explore this, we analyzed Cell Painting and L1000 data for 106 compounds across three cell lines from osteoblast, lung, and breast tumors (U2OS, A549, and MCF7) at three time points (6h, 24h, 48h) using a 10µM concentration. We found significant time and cell line effects in Cell Painting data, with less pronounced effects in transcriptomics. Using Weighted Gene Co-expression Network Analysis (WGCNA) and enrichment analysis, we identified connections between cell morphology and gene deregulation for chemicals with similar biological effects (e.g., HDAC and CDK inhibitors). These findings suggest that while Cell Painting shows distinct patterns, both technologies offer complementary insights into compound-induced cellular changes, enhancing drug discovery and chemical risk assessment.

## Introduction

In many successful drug discovery programs, combining target knowledge with functional cellular assays is essential for identifying potential drug candidates^1^. Furthermore, to uncover the molecular mode of action (MoA) of a chemical, integrating multi-omics data, such as transcriptomics, proteomics, epigenomics, and metabolomics, has been explored using advanced technologies^2,3^. However, the use of these technologies remains relatively expensive for high-throughput studies.

For the past few years, there has been a revival of phenotypic screening in drug discovery, particularly for understanding disease biology that can be modulated by small molecules^1,4^. More recently, the same approaches have been applied in safety profiling to better understand the potential of candidate drugs to induce adverse outcomes. Advances in various technologies for cell-based phenotypic screening, including the development of induced pluripotent stem (iPS) cell technologies, gene-editing tools such as CRISPR-Cas, and advanced imaging assays, have become integral components of the pharmaceutical industry’s toolkit. These tools facilitate the systematic determination of cellular effects caused by chemicals^5,6^. More recently, such bioactivity data have also been shown to be useful for de novo compound design. This involves algorithmically generating novel chemical compounds with specific desirable properties or biological activities, with promising applications in hit discovery^7,8^.

Phenotypic screening strategies stand out for its ability to simultaneously measure multiple morphological parameters up to the single-cell level. Cell Painting, a widely used high-content screening (HCS) technique, captures cellular morphological changes and translates them into thousands of features, effectively assessing responses to genetic modifications, environmental stressors, and small molecule treatments^9^.

In contrast to target-based approaches, phenotypic screening does not require knowledge of the molecular target perturbed by a stressor, making it challenging to translate molecular mechanisms of action in disease-relevant cell systems using only HCS. However, studies have shown that cell morphology data can complement gene expression or chemical structure information. For example, Seal et al. found that Cell Painting readouts can cluster various modes of action of mitochondrial toxicants^10^. Schneidewind et al. used morphological profiling to group biosimilar compounds with shared modes of action (MoA)^11^. Cell Painting fingerprints have already been used to associate groups of chemicals with biological pathways, targets or diseases^12,13^. Combined with gene expression profiling or structural information, Cell Painting has also proven to be very promising for predicting various toxicity endpoints, such as drug-induced liver injury^14,15^, acute toxicity^16^, and cardiotoxicity^17^.

The impact of Cell Painting over the past 10 years in life sciences and drug discovery was reported in a recent comprehensive review^18^. Overall, numerous studies agree that combining multiple screening technologies, such as Cell Painting and transcriptomics, is key to predicting and understanding the modes of action of chemical compounds for both pharmacological and toxicological properties^19–21^. The potential of a multimodal profile has been explored multiple times, either through a comparative analysis of phenotypic and transcriptomic profiles^22^, or by integrating the two types of profiles into a biological network^13^. Way’s study, based on Cell Painting and gene expression data from A549 cells treated with a large set of compounds, concluded that morphological profiles capture more cellular states than gene expression profiles^22^.

Although promising, these studies combining phenotypic screening and gene expression profiling were reported on a single cell line. However, it has been demonstrated that the choice of cell line is crucial, whether in Cell Painting^23,24^ or in transcriptomics^25^. We believe the field lacks a systematic study assessing the specificity and complementarity of gene expression and morphological changes over time across different cell lines in response to chemical perturbations.

Here, we gathered recently released gene expression and Cell Painting data for a collection of 106 chemicals^26,27^. Chemicals were tested on three cell lines originating from osteoblast (U2OS), lung (A549), and breast (MCF7) tumors at 10µM, measured at 6h, 24h, and 48h. Our analysis showed clear cell line and time effects in Cell Painting data, which were less obvious in gene expression data. These results suggest specific cell morphology patterns dependent on cell line and time point. Additionally, we assessed the relationship between Cell Painting and gene expression using Weighted Gene Co-expression Network Analysis (WGCNA). WGCNA has been widely used to identify and cluster genes^28–30^ deregulated in the same manner, but it has never been applied to Cell Painting data. By implementing this method on both data types and combining it with customized enrichment analysis, we observed that molecules with similar modes of action, notably HDAC and CDK inhibitors, could be clustered together in both Cell Painting and gene expression data, establishing a relationship between the readouts from both technologies.

Overall, this analysis suggests that cell morphology profiles are more specific to duration of exposure and cell lines than gene expression profiles under the tested conditions. However, mechanisms of action related to gene expression changes can also be reflected in cell morphology perturbations, offering insights for cell profiling in response to chemicals.

## Results

### Visualizing impact of both cell line and time effects on cell morphology and gene expression

Using three cell lines (U2OS, A549, MCF7) exposed to a set of 106 chemicals at three time points (6h, 24h, 48h), we first analyzed the readouts of Cell Painting (CP) and LINCS (L1000) in the aim to compare the synergy or diversity in cell morphology and transcriptomic modulation induced by the chemical stressors across all the tested conditions. We started by generating two Uniform manifold Approximations and Projections (UMAPs) of the profiles corresponding to the 106 compounds tested in each cell line and duration of exposure, in either CP (Figure 1A) or L1000 (Figure 1B). For both CP and L1000, the UMAPs did not reveal any discernible impact of either time or cell line on the controls (DMSO) (Figure 1). DMSO of every condition coinciding in the center of the projections suggested that the observed clustering, particularly in CP, was not due to batch effects.

**Figure 1.**
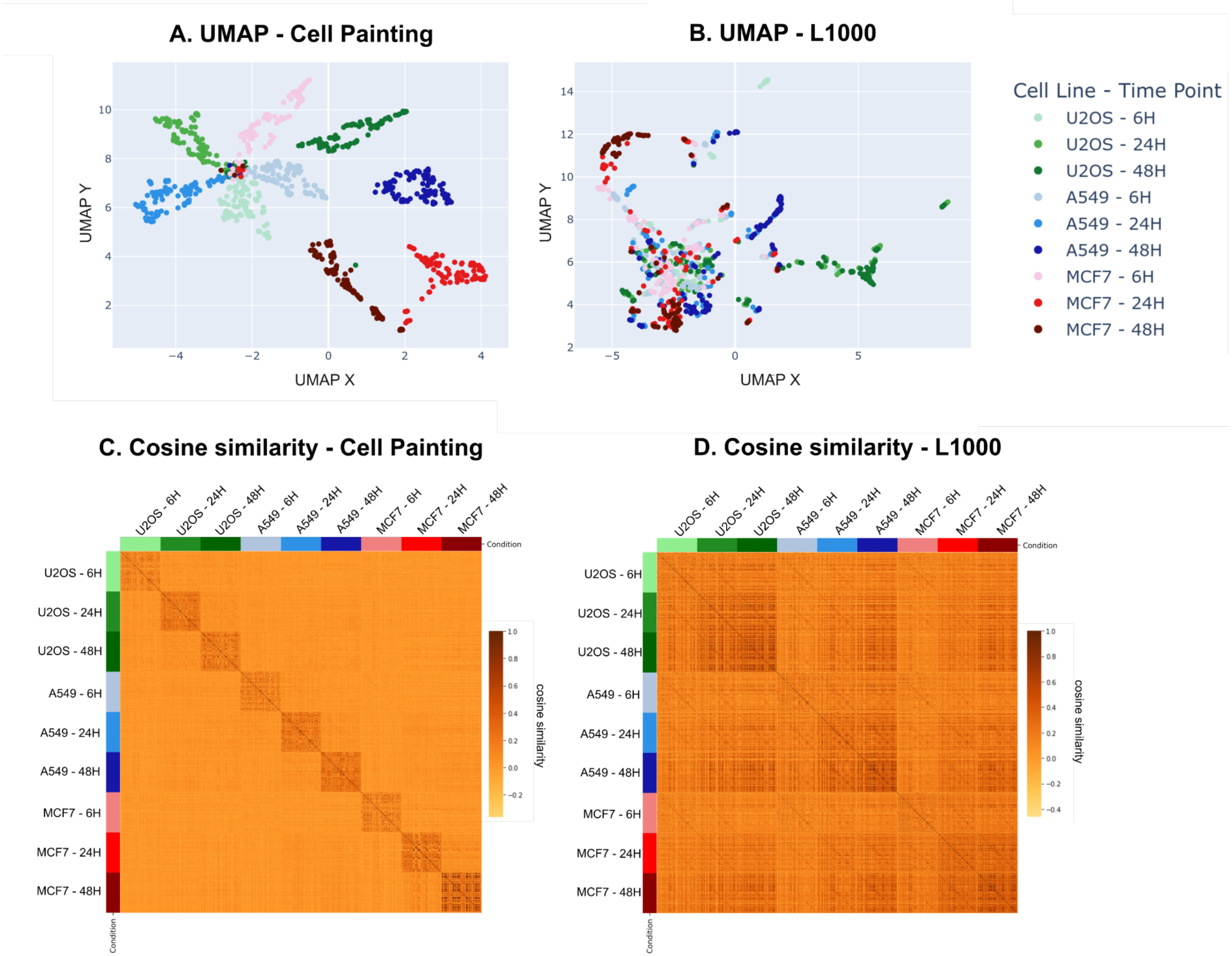
Comparison of the morphological and transcriptomic responses between cell lines and over duration of exposure. **A** and **B** are UMAPs where each point represents the profile of a compound tested in either CP **(A)** or L1000 **(B)** within a specific cell line and time point. UMAPs are color coded by the different cell lines: U2OS in green, A549 in blue, and MCF7 in red. The colors transition from lighter to darker as the exposure time increases, going from 6h to 48h. Heatmaps in **C** and **D** show the cosine similarity between the compounds tested in every condition of cell line and time in either CP **(C)** or L1000 **(D)**. The profiles were grouped by condition. The associated color code, on the side of the heatmap, is the same as the one used on the UMAPs.

Figure 1A showed that the CP profiles clustered exclusively based on cell line and exposure time, independent of the chemical tested. We also observed that the three clusters corresponding to the compounds tested at 6h in the three cell lines were located closer to the control at the center of the projection, while the three clusters corresponding to the compounds tested at 48h in the three cell lines were situated on the right side of the projection. Interestingly, after 24 hours of exposure, we noticed that the two clusters for the U2OS and A549 cell lines were on the left of the projection, while the one corresponding to MCF7 was on the far right. Nevertheless, aside from an exception at 24 hours for MCF7, a trend seemed to emerge, with the phenotypic spaces between cell lines tending to converge for a given exposure time.

The biological spaces associated with the transcriptomic responses were different (Figure 1B). The UMAP did not allow for the isolation of distinct transcriptomic responses of the compounds between the cell lines after 6 hours of exposure, as we observed a single cluster (Figure 1B). However, it also revealed that, particularly at 48h, transcriptomics profiles tended to cluster mode according to the cell line in which the compounds have been tested. This highlighted the emergence, starting from 24h, of a cell line effect on the transcriptomic response between U2OS, A549 and MCF7 cell lines, which seemed to be correlated with the increase of the duration of exposure.

In order to confirm the observations from the UMAPs, we calculated the cosine similarity between the profiles in each condition for CP and L1000. In accordance with UMAPs, similarity between morphological profiles was higher between different compounds tested in the same conditions of time and cell line than between same compounds but tested in different conditions (Figure 1C). The study of similarity between L1000 profiles depicted the prevalence of a cell line effect on the compound effect, showing at 48h a higher cosine similarity between compounds within the same cell line compared to the same compounds tested at shorter exposure times but in different cell lines (Figure 1D).

Overall, the UMAPs showed a clear distinction in cell responses at the morphological level compared to the transcriptomic level, where the modulation after chemical exposure is more subtle and may be related to the deregulation of a few genes.

### Identifying phenotypic and transcriptomic biological spaces related to HDAC and CDK inhibitory compounds

We were interested in the behavior in both CP and L1000 of compounds sharing similar modes of action (MoA). Therefore, we annotated the compounds with their known MoA. Among the most represented MoA, 10 compounds were identified as histone deacetylase (HDAC) inhibitors (panobinostat, vorinostat, resminostat, PCI-24781, JNJ-26481585, belinostat, entinostat, romidepsin, tacedinaline and dacinostat), and 9 compounds as cyclin-dependent kinase (CDK) inhibitors (R547, PHA-793887, AZD5438, dinaciclib, AT-7519, alvocidib, JNJ-7706621, roscovitine and palbociclib). These modes of action being the most represented in our dataset, we chose to focus on those two. For both CP features and L1000 genes and by cell lines, we created UMAP gathering all the compounds tested at the three time points. Figure 2 shows the position of HDAC and CDK inhibitors among the other compounds at 6h, 24h and 48h, in the A549 cell line, in the phenotypic and transcriptomic spaces.

**Figure 2.**
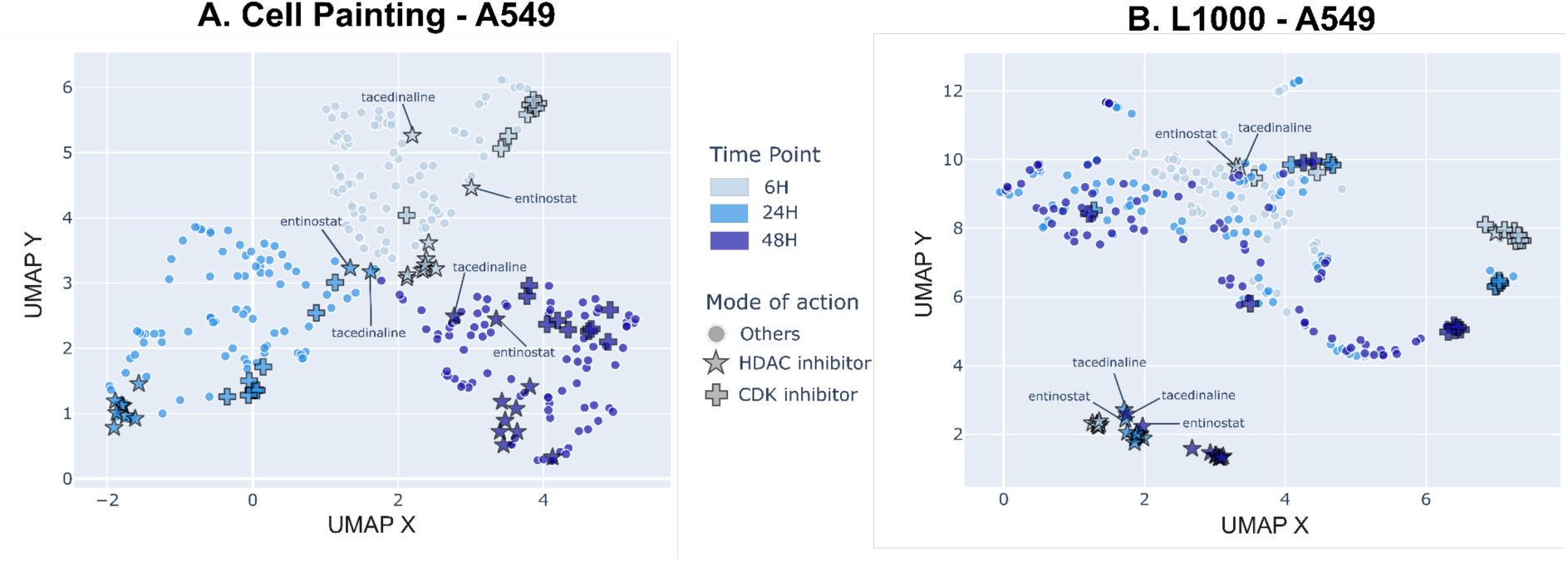
Comparison over time of the phenotypic and transcriptomic signatures of HDAC and CDK inhibitors in the A549 cell line. Each point represents a compound tested in either CP **(A)** or L1000 **(B)** on A549 cells. The colors transition from lighter to darker as the exposure time increases, going from 6h to 48h. HDAC inhibitors and CDK inhibitors are respectively symbolized by stars and crosses outlined in black, and the other compounds by circles. Points corresponding to entinostat and tacedinaline were specifically labeled.

It came out from Figure 2A that inside each of the three main clusters, corresponding to the compound’s morphological signatures at 6h (light blue), 24h (blue) and 48h (dark blue) in the A549 cell line, 2 subclusters of respectively HDAC and CDK inhibitors (respectively stars and crosses) could be observed. With L1000, the spaces associated with the duration of exposure were less perceptible than with CP, especially at 24h and 48h that tended to overlap (Figure 2B). The majority of the HDAC inhibitors were grouped at the bottom of the UMAP, suggesting a transcriptomic signature of the HDAC inhibitors in the A549 cell line independent of the time, with the exception of entinostat and tacedinaline. We noticed the particular case of the over time signature of entinostat and tacedinaline in CP too. At 6 hours, these two compounds already had a strong signature, more distant from those of the other HDAC inhibitors, which remained close to the controls at the center of the map (Figure 2A). However, at 24 hours and 48 hours, the perturbations of entinostat and tacedinaline seemed less important (closer to the center of the map and the controls), while signatures of the other HDACs were highly differentiated at the extremes of the clusters. It appears that entinostat and tacedinaline have effects on A549 cells with both technologies, at shorter exposure times compared to the other HDAC inhibitors.

Regarding the CDK inhibitor signatures, they were not as pronounced as the HDAC inhibitors ones, as we encountered 2 subgroups of compounds (Figure 2B). The first subgroup, in the central part of the map, regrouped mostly roscovitine and palbociclib at 6h, 24h, 48h, and PHA-793887 and JNJ-7706621 at 24h and 48h. The second one was at the extreme right side of the map with the rest of the CDK inhibitors, sorted by duration of exposure.

Same UMAPs for the U2OS and MCF7 cell lines are also available in Figures S1 and S2. The clustering obtained for the U2OS and MCF7 cell lines was generally similar (Figures S1 and S2). Whether the biological spaces related to cell lines and time points were distinctly separated as seen in CP or more blended as observed in transcriptomics, the HDAC and CDK inhibitors consistently clustered within specific subspaces. This confirms that the MoA of these chemicals can be detected at both the transcriptomic and cell morphology levels.

### Assessing the correlation between modules of morphological features and modules of genes through customized enrichment

Understanding the morphological perturbations induced by chemicals, especially from a mechanistic perspective, remains one of the principal challenges in CP. We hypothesized that compounds that perturb a set of morphological features in a similar manner could also affect the same set of genes. To answer this question, we developed a three-step protocol. First, we implemented a WGCNA approach allowing to respectively group co-expressed genes and co-perturbed features into modules. Then, as the modules are independent of the compounds, we computed a customized enrichment score to link each compound to each module, measuring the extent to which a compound deregulates each module. In the final step, for a given condition, we clustered the compounds based on their enrichment score for the gene modules, and in parallel, we clustered these same compounds based on their enrichment score for the morphological feature modules. Using a tanglegram representation, we compared these two clustering and underlined a correlation between the transcriptomic and phenotypic spaces by measuring the impact of each compound on each module of genes/features across different time exposure conditions and cell lines.

As an example, Figure 3 depicts the correspondence at the 6h time point in each cell line between compounds that tend to enrich the same gene modules and those that tend to enrich the same morphological feature modules. The tanglegram visualization allowed us to link the leaves (representing compounds) of two dendrograms. Thus, a group of parallel links indicates a cluster of compounds that is found in both CP and L1000, and consequently a potential relationship between the modules of genes and features enriched by the compounds of this cluster. At first glance and considering the tanglegrams of each cell line individually, we observed several clusters of compounds which look similar between CP and L1000 at 6h (Figure 3).

**Figure 3.**
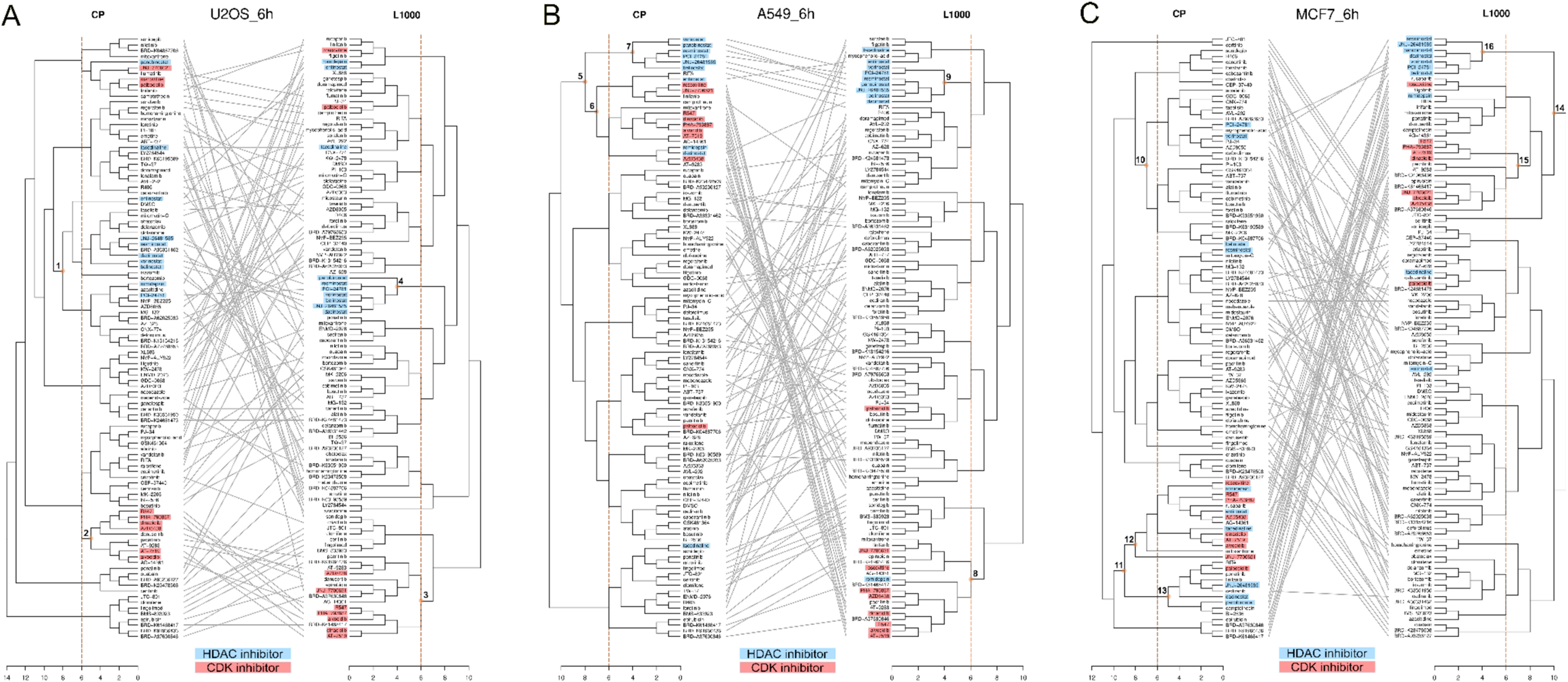
Tanglegrams comparing the hierarchical clustering of compounds at the 6h time point in the U2OS, A549, and MCF7 cell lines. Individual dendrograms were computed separately using distance matrices calculated with Euclidean distance and clustered with the ward.D2 method. A distance scale is located at the bottom of each dendrogram. The left dendrograms represent the clustering of compounds based on their customized enrichment scores for modules of CP features, while the right dendrograms represent clustering based on the same enrichment scores for modules of L1000 genes. Then, tanglegram visualization enables comparison between CP and L1000 dendrograms, where each leaf is a compound tested at 6h, for U2OS **(A)**, A549 **(B)** and MCF7 **(C)**. HDAC inhibitors are highlighted in blue, and CDK inhibitors in red. To facilitate the description of the tanglegrams, the most interesting branches were marked and numbered. The vertical orange line at a clustering distance of 6 was used to define HDAC and CDK inhibitor cores as described in the Methods.

Given the pharmacological annotations, we highlighted either in blue or red, compounds that we know HDAC or CDK inhibitors. For each of the studied cell lines, the clustering of compounds based on the enriched L1000 gene modules revealed that HDAC inhibitors and CDK inhibitors tended to be grouped within the same clusters, respectively. A group of HDAC inhibitors, consisting of panobinostat, resminostat, PCI-24781, vorinostat, belinostat, JNJ-26481585, and dacinostat, was consistently found in the same cluster on the L1000 side in U2OS, A549, and MCF7 (Figure 3, branches no. 4, 9, 16). On the CP side, this same group was found in the A549 cell line (Figure 3B, branch no. 7), and a similar group was found for U2OS (Figure 3A, branch no. 1). However, HDAC inhibitors likely impacted cell morphology more diversely in MCF7, as they were not clustered together (Figure 3C, branches no. 10, 12, and 13). Afterwards, in all three cell lines at 6 hours, there was a clear link between the morphological and transcriptomic effects of CDK inhibitors, as they tended to cluster together when considering both the enrichment of feature modules and gene modules (Figure 3, branches no. 2, 3, 6, 8, 12, and 15). The least correlation was observed between CP and L1000 for the behavior of CDK inhibitors in U2OS. Interestingly, we detected that among the CDK inhibitors, roscovitine and palbociclib were separated from the main CDK inhibitor cluster in the three cell lines at 6 hours (Figure 3), whether in CP or with L1000. However, roscovitine and palbociclib are first- and third-generation CDK inhibitors, respectively, while all the others are second-generation, and thus have different mechanisms of action^31^. These differences in mechanisms for the same type of compound resulted in different morphological and transcriptomic perturbations as well.

Finally, we noticed that in MCF7 cells, HDAC inhibitors seemed to have morphological and transcriptomic effects similar to those of CDK inhibitors, as they were found in the same part of the dendrogram in both CP and L1000 (Figure 3C, branches no. 11, 14). This was also the case morphologically in A549 cells (Figure 3B, branch no. 5). It could be explained by the fact that HDAC and CDK inhibitors are both types of anticancer drugs^31,32^, mostly inducing cell cycle arrests and cell death.

Tanglegrams were also generated for the 48h time point (Figure S3). Similar to the 6h analysis, clusters for HDAC and CDK were observed. No new clusters of interdependent compounds between CP and L1000, specific to longer exposure durations, were identified.

Overall, the module enrichment analysis revealed that compounds sharing the same pharmacological role can exhibit both common morphological and transcriptomic signatures.

### Creating a bipartite signature of HDAC inhibitor and CDK inhibitor compounds

The ultimate goal would be to deduce a typical profile combining genes and morphological features for a given type of drug or chemical. We focused on the cores of HDAC and CDK inhibitors defined in Methods. From the calculated enrichment scores between each compound and WGCNA module of genes or cell morphology features, we performed Mann-Whitney tests for each cell line and time condition in the aim to identify gene modules and features that were significantly more enriched by the HDAC and CDK core compounds than by the others. Selected gene modules and CP modules per condition are available in Table S1.

For HDAC and CDK inhibitors and for each condition, the combination of both CP and L1000 representative modules could compose a typical profile associating the morphological features and genes that are simultaneously impacted by these types of drugs. In order to understand more precisely each typical profile and how HDAC or CDK inhibitors are perturbing the cells in each condition, we first performed gene enrichment on the HDAC or CDK inhibitors representative modules. The analyses revealed biological pathways disrupted by the genes in the representative modules of HDAC or CDK inhibitors in each cell line, both at 6 hours and at 48 hours. Among the biological pathways disturbed by the HDAC inhibitors, we notably found the apoptosis signaling pathway, the kinase activity, the mitosis and cell cycle, the endoplasmic reticulum (ER) response to stress, or the process of endocytosis. Some were expected, as HDAC inhibitors are for example known to induce apoptosis and cell cycle arrests in cancer cells or to modulate transcription^33^. Simultaneously, gene enrichment also highlighted that cellular components such as the nucleus, granules and vesicles managed by the golgi apparatus, and ER were perturbed by HDAC inhibitors. This allows us to propose specific links with the CP feature modules. Cell Painting features can be categorized by channels representing cellular compartments^34^. For each condition, CP modules representative of HDAC inhibitors contained features belonging to the nucleus, where DNA replication takes place during the cell cycle or some protein phosphorylation during transcription, the ER, or the golgi apparatus (Table S1).

In the case of CDK inhibitors at the 6h time point, still the apoptosis signaling pathway, the kinase activity, the mitosis and cell cycle, the ER response to stress, but also the immune response of the cell, in the mitochondrial catabolic process or in the cytoskeleton and fiber assembly were associated with CDK inhibitors. Once again, these results were consistent with the known mechanisms of action of CDK inhibitors, namely modulation of the cell cycle, induction of apoptosis, and enhancement of the immune response^31^. At 48h, genes of the CDK inhibitors representative modules mostly disturbed the cell cycle, notably the mitosis, the kinase activity, the immune response or the apoptosis. Except for MCF7 at 48h, corresponding CP channels and so cellular compartments associated with all of these pathways were also represented by the features of the CDK inhibitors representative modules (Table S1). After 48 hours of exposure to CDK inhibitors, the effects on the cell as a whole could be more pronounced. Consequently, it would become more challenging to identify specific disruptions in a particular cellular compartment.

### Comparing representative modules of HDAC inhibitors or CDK inhibitors over time and between cell lines

Results detailed above showed that the biological pathways and cellular compartments impacted with HDAC or CDK inhibitors were similar over cell lines. We wanted to see if the paired morphological and transcriptomic signature combining HDAC or CDK inhibitors representative modules was stable over time and between U2OS, A549 and MCF7. We reused the significantly representative modules indicative of the core of HDAC inhibitors or CDK inhibitors that we previously isolated, and compared the morphological features and genes contained in these modules between the cell lines and at 6 hours and 48 hours. Figure 4 showed for HDAC inhibitors and CDK inhibitors the numbers of shared features and genes, all representative modules combined, between conditions.

**Figure 4.**
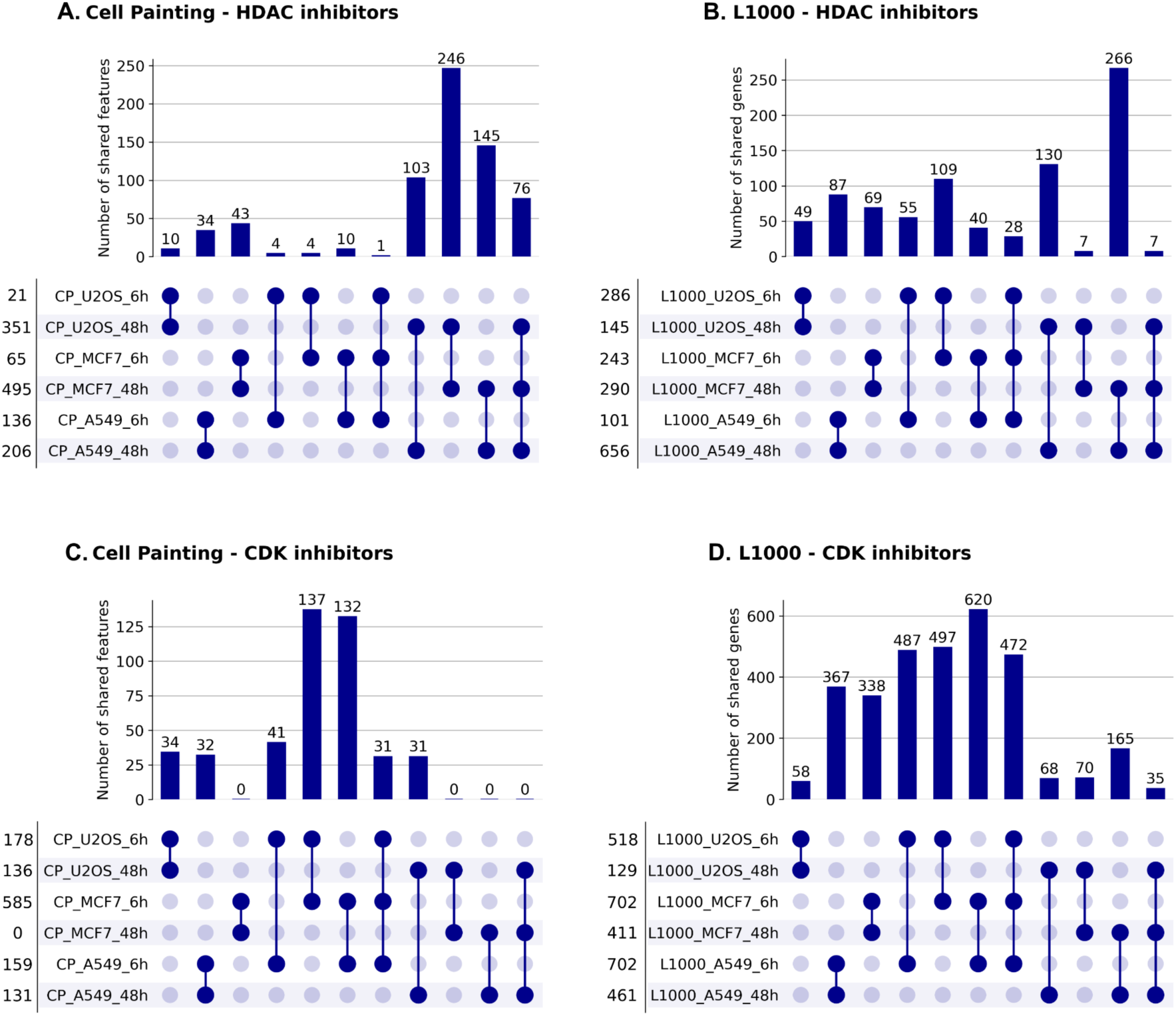
Overlap between cell lines and over time of features and genes included in HDAC and CDK inhibitors-discriminative modules. Each row below the upset plot represents a condition (cell line and duration of exposure). The number to the left of each condition is the number of morphological features or genes belonging to the modules that were significantly enriched by the core HDAC inhibitors or the core CDK inhibitors (all modules representative modules combined). Above, bars represent the number of shared morphological features or genes between different combinations of conditions studying cell line or time effects. **(A)** Number of features shared between conditions for HDAC inhibitors. **(B)** Number of genes shared between conditions for HDAC inhibitors. **(C)** Number of features shared between conditions for CDK inhibitors. **(D)** Number of genes shared between conditions for CDK inhibitors.

As mentioned before, HDAC inhibitors-discriminative modules of features contained in each cell line at 48 hours more features than at 6h. Accordingly, we also noticed that 76 features were shared by the HDAC inhibitors-discriminative modules of morphological features between the 3 cell lines at 48 hours, against only 1 (that concerned mitochondria) at 6h (Figure 4A). Among the 76 shared features between the 3 cell lines at 48h, 13 were associated with the nucleus, 22 with RNA, 14 with ER, 15 with mitochondria, and 17 with AGP (Table S2). On the contrary, at the molecular level, only 7 genes were present in the HDAC inhibitors-discriminative modules of genes of each cell line at 48h, whereas they were 28 at 6h (Figure 4B). Among these 7 genes, we notably noticed *WIPF2*, *PIK3C3* and *HDAC6* (Table S2). *WIPF2* encodes a protein involved in the organization of the actin cytoskeleton^35^, and *PIK3C3* results in a protein playing a role in the functional of early to late endosome but also in the ER traffic^36^. All these mechanisms are directly related to the previously highlighted biological pathways and cellular components that are perturbed by HDAC inhibitors. Finally, *HDAC6* is known to be a biological target of the HDAC inhibitors. It participates in the regulation of the transcription, the cell cycle, and its inhibition can damage actin cytoskeleton-dependent cell migration^37^. Moreover, *HDAC6* subcellular locations are plasma membrane, cytoskeleton, cytoplasm, endosome and nucleus^37^. *WIPF2*, *PIK3C3* and *HDAC6* could be interesting genes in the isolation of a shared transcriptomic signature of HDAC inhibitors at 48h between U2OS, A549 and MCF7.

Considering cell lines separately, the preservation of the morphological and transcriptomic signature over time of HDAC inhibitors would be conceivable. Indeed, in each cell line, HDAC inhibitors-representative modules at 6 and 48h systematically shared some features (U2OS: 10, A549: 34, MCF7: 43) (Figure 4A) and genes (U2OS: 49, A549: 87, MCF7: 69) (Figure 4B). For HDAC inhibitors in each cell line, there are overlapping patterns in both morphological and transcriptomic signatures over time.

Regarding the CDK inhibitors, each cell line, and particularly MCF7 were responsive in both morphological and transcriptomic aspects, with 31 features (Figure 4C) and 472 genes (Figure 4D) shared by the CDK inhibitors-discriminative modules of the three cell lines. Among the 31 cell morphology features, 7 referred to the nucleus, 6 to the nucleolus and cytoplasmic RNA, 4 to the ER, 7 to the mitochondria and 4 to others (Table S2). Moreover, the 472 genes representative of the common signature of CDK inhibitors in U2OS, A549 and MCF7 at 6h included *CDK4*, *CDKN2A* and *E2F2*. *CDK4* encodes for CD4 protein, which usually complexes with cyclin D, enabling the cell to enter in phase S of mitosis, through the activation of protein from the E2F family^31,38^. *CDK4* is a well-known target of some CDK inhibitors. *CDKN2A* codes for protein p16, which can fix CDK4 and prevents it from interacting with cyclin D, thus blocking the entry in phase S^39^. This suggests the potential for a common signature of CDK inhibitors across the three cell lines at 6 hours.

However, the idea of a paired morphological and transcriptomic signature over time common to U2OS, A549 and MCF7 was made complicated by the morphological response of the MCF7 cells at 48h. CDK inhibitors-discriminative modules of features in MCF7 included at 6h 585 features out of the 732 features of the dataset, but none at 48h (Figure 4C). However, an over time morphological and transcriptomic signature of CDK inhibitors would be possible for U2OS, including 34 genes and 58 features, and for A549, with 32 features and 367 genes (Figure 4C, 4D). Moreover, this signature could be common to these 2 cell lines, as CDK inhibitors-discriminative modules of features shared 41 morphological features at 6h and 31 at 48h (Figure 4C), and CDK inhibitors-discriminative modules of genes shared 487 genes at 6h, and 68 at 48h (Figure 4D).

## Discussion

Understanding and predicting how drugs interact with biological systems is challenging due to the complexity of multiple interacting pathways and networks. Drugs can have various targets and off-target effects, making their overall impact difficult to predict, especially at the whole organism level. Making use of high-content screening assays offers promise in addressing this major challenge in drug discovery and chemical risk assessment. Various factors must be considered to elucidate the mechanisms by which a chemical impacts a biological system. By leveraging advanced technologies in cell-based phenotypic screening (e.g., Cell Painting) alongside high-throughput transcriptomic profiling, we assessed the differences and complementarity of responses at three time points and in three cell lines for a set of 106 diverse chemicals tested at one concentration. Although previous studies reported that a same compound could induce a different phenotypic profile^24^ or transcriptomic profile depending on the compound MoA^25^ across cell lines. We pointed out distinct phenotypic spaces specific to U2OS, A549 and MCF7. Interestingly, these morphological spaces evolved over time and were specific to each of the three cell lines tested. In contrast, the biological spaces overlapped more significantly when examining transcriptomic responses in the same conditions. Distinct transcriptomic responses for each cell line became more apparent with longer exposure times. This may be explained by the fact that L1000 measures the mRNA levels of 978 transcripts representing highly ubiquitous genes^27^. Using a whole transcriptome dataset would be advantageous for capturing transcripts that are more specific to each cell line. Nevertheless, HDAC and CDK inhibitors showed both homogeneous morphological and transcriptomic profiles when considering each cell line individually. Moreover, the different time responses of HDAC and CDK inhibitors were generally well captured by Cell Painting, but also more slightly in L1000.

Although the complementarity and differences between cell morphology and gene expression profiling in terms of reproducibility, endpoint prediction, and mode of action analysis have already been studied^13,22,40^, these studies did not allow for a direct connection between genes and morphological features. Having demonstrated distinct patterns for HDAC and CDK inhibitors using both technologies across three cell lines and at different time points, we aimed to identify useful overlaps between cell morphology and gene expression readouts that could be valuable in a multi-modal profile.

Our approach involved a combination of WGCNA with a customized enrichment score to associate sets of genes and cell morphology features with each compound individually. We demonstrated that HDAC inhibitors and CDK inhibitors enriched the same sets of genes and the same sets of Cell Painting features, indicating that common gene and cell morphology profiles could be defined for the same types of compounds using both technologies, and offering a better characterization of their mechanisms of action through the identification of both key genes and morphological features. By coupling both profiles, we deduced a gene-morphology relationship that can also be preserved between several cell lines or over time. This is a promising result as establishing a linkage between perturbations at the molecular and cell morphology levels is highly challenging and faces significant bottlenecks^22,41^.

However, our study also indicates rooms for improvement. Firstly, the WGCNA methodology was unsuccessful in creating modules of cell morphology features in A549 and MCF7 cell lines at the 24h time point. Although the WGCNA protocol has already been adapted to other technologies^42^, the high correlation among Cell Painting features may be a limiting factor in our protocol, complicating the identification of networks between these features. Furthermore, the study utilized carcinoma cell lines, which could be relevant for examining the effects of HDAC and CDK inhibitors, commonly employed in cancer therapy^37,43,44^. A similar analysis using primary human cells or induced pluripotent stem cell (iPSC)-derived cells, which are more representative of human physiology^45^, would be more suitable for drug discovery. Finally, our analysis has been done on 1 concentration. The recent study by Way et al., which assesses the Cell Painting and L1000 perturbations of compounds across 6 doses in A549 cells^22^, would be interesting to investigate with our approach.

Overall, this study demonstrates the differences and complementarity of Cell Painting and L1000 technologies using different cell lines and time points to detect perturbations caused by chemicals. Moreover, genes and cell morphology profiles can be characterized from both readouts and combined into a unique profile that can be related to groups of chemicals sharing a similar mode of action. Further collaborative efforts are underway, including supplementary experiments that will expand profiling beyond phenotypic and transcriptomic analyses to include additional types such as proteomics (OASIS) (https://oasisconsortium.org/) and metabolomics (RiskHunt3R) (https://www.risk-hunt3r.eu/). Integrating such multimodal profiles and exploring these data through innovative data science technologies could enhance the power and resolution of chemical profiling in biology. This approach could significantly advance phenotypic drug discovery and chemical risk assessment.

## Acknowledgement

This work was supported by the project RISK-HUNT3R: RISK assessment of chemicals integrating HUman centric Next generation Testing strategies promoting the 3Rs. RISK-HUNT3R has received funding from the European Union’s Horizon 2020 research and innovation program under grant agreement No 964537. This work is also supported by Bayer Crop Science through GIE AIFOR PhD Cifre funding (2021-0755).

## Author contributions

VL: conceptualization, data processing, data curation, data analysis, methodology, visualization, writing. DR: conceptualization, funding acquisition, writing and editing. OT: conceptualization, funding acquisition, writing.

## Supplementary information

Document S1. Figures S1-3. (PDF)

Table S1. Details of WGCNA modules for Cell Painting and L1000. Each sheet corresponds to a condition. WGCNA modules are named by default by color. We highlighted blue HDAC inhibitor representative modules only, red CDK inhibitor representative modules only, and purple modules that were representative of both HDAC and CDK inhibitors. (XLSX)

Table S2. Details of overlaps between HDAC inhibitors and CDK inhibitors representative modules. First two sheets describe overlaps between HDAC inhibitor representative modules for Cell Painting and L1000. Last two sheets describe overlaps between CDK inhibitor representative modules for Cell Painting and L1000. (XLSX)

Table S3. Details of the 106 selected compounds and annotations. (XLSX)

## STAR Methods

### Key resources table

| REAGENT or RESOURCE | SOURCE | IDENTIFIER |
| --- | --- | --- |
| **Software and Algorithms** |  |  |
| Python | Rossum et al.<sup>48</sup> | v3.7.12 and v3.9.16 |
| R | R Core Team<sup>53</sup> | v4.3.2 |
| cytomniner-eval | https://github.com/broadinstitute/grit-benchmark | v0.1 |
| umap-learn | McInnes et al.<sup>50</sup> | v0.5.1 |
| seaborn | Waskom et al.<sup>51</sup> | v0.12.2 |
| WGCNA | Langfelder et al.<sup>52</sup> | v1.72-5 |
| EnrichIntersect | Zhao et al.<sup>54</sup> | v0.6 |
| dendextend | Galili et al.<sup>55</sup> | v1.17.1 |
| upsetplot | Lex et al.<sup>56</sup> | v0.9.0 |
| GSEAPy | Fang et al.<sup>57</sup> | v1.1.0 |
| **Others** |  |  |
| Normalized Cell Painting profiles at the well-level | Natoli et al.<sup>26</sup> | https://doi.org/10.5281/zenodo.5008187 |
| Cell Painting processing workflow | Natoli et al.<sup>26</sup> | https://github.com/broadinstitute/lincs-cell-painting |
| L1000 profiles at the level 5 (beta version of 2020) | L1000 assay<sup>27</sup> | https://clue.io/releases/data-dashboard |
| Compound annotations | Natoli et al.<sup>26</sup> | https://github.com/broadinstitute/lincs-cell-painting |

### Code and data availability

- This paper analyzes existing, publicly available data. These datasets and their source are listed in the key resources table.
- The original code can be shared by the lead contact upon request.
- Any additional information is available from the lead contact upon request.

### Method details

#### Cell Painting data

The Cell Painting (CP) data, which represents morphological profiles, were obtained from a Zenodo archive provided by Natoli et al.^26^. We specifically selected the second batch of data. When considering the perturbation identifier, we compiled morphological profiles for 331 compounds at 3 time points (6h, 24h, and 48h), at a concentration of 10µM, and across 3 cell lines: U2OS (human osteosarcoma), A549 (human lung carcinoma), and MCF7 (breast cancer). The number of replicates per compound ranged from 3 to 35 depending on the conditions. The profiles were obtained at level 4b, indicating that single-cell profiles were extracted from the Cell Painting images, aggregated to the well level, annotated with metadata, normalized over controls (DMSO) using the “mad_robustize” method, and finally spherized over DMSO’s. The spherizing process aims to correct batch effects^46,47^. From the original set of 2198 features, a feature selection step was performed before the spherize transformation, involving the removal of features with low variance, high correlation, excessive missing values, or outliers. This process resulted in the selection of 732 features. The entire data processing workflow is described in detail in the publicly available GitHub repository at https://github.com/broadinstitute/lincs-cell-painting^26^.

#### Categorization of the Cell Painting morphological features

As described by Bray et al., morphological features were extracted with CellProfiler and can be sorted regarding the stained cell organelles^34^: nucleus (DNA channel), nucleolus and cytoplasmic RNA (RNA channel), F-actin cytoskeleton, Golgi, and plasma membrane (AGP channel), endoplasmic reticulum (ER), mitochondria (Mito), and others, which concern features dealing with the general size or shape of the cells. For correlation features that provide a measurement based on two channels, we have decided to consider them as a feature of each of these two channels.

#### L1000 data

The gene expression profiles were obtained from the L1000 assay, developed by the NIH LINCS Consortium and Subramanian et al.^27^. The L1000 assay provides gene expression profiles for 978 landmark genes, processed at different levels ranging from raw fluorescent intensities (level 1) to differential gene expression signatures (Z-scores) where biological replicates are aggregated based on their average correlation (level 5). We obtained L1000 profiles (beta version of 2020) from a dashboard provided by the L1000 assay, accessible at https://clue.io/releases/data-dashboard. The profiles were collected at level 5, encompassing data from 3 cell lines (U2OS, A549, MCF7), at 3 time points (6h, 24h, and 48h), and for a concentration of 10µM. Out of the 331 compounds in the Cell Painting data, identified by perturbation identifier, we successfully retrieved L1000 profiles for at least one time point for 322 compounds in U2OS, and 331 compounds in both A549 and MCF7, with between 1 and 93 replicates depending on the conditions.

#### Selection of strong and reproducible Cell Painting and L1000 profiles

In the aim to consider only robust CP profiles with a high confidence of reproducibility, we used the grit score metric, designed by the Broad Institute to filter Cell Painting profiles according to their reproducibility and phenotypic strength. It takes into account replicates reproducibility and distance between compounds and controls. We computed the grit score for each level 4b Cell Painting profiles using the implementation available in the cytomniner-eval Python package (v0.1, Python v3.7.12^48^) of the Broad Institute (https://github.com/broadinstitute/grit-benchmark). Considering that a grit score can be assimilated to a Z-score, we excluded compounds for which the average grit score of replicates was less than two^49^.

We operated a similar selection on L1000 profiles using two metrics, the Replicate Correlation Coefficient (CC) and the Signature Strength (SS), detailed in the supplementary information in Sabramanian, et al.^27^. CC and SS scores were provided with the L1000 profiles. By filtering the compounds based on the mean CC and SS values of the replicates (replicate mean CC>=2 and replicate mean SS>=200), we selected compounds considered as reproducible and robust in L1000.

These two protocols allowed the selection of 106 compounds (DMSO included), that encompassed the defined criteria of grit score, CC and SS in every cell line and time point. Perturbation replicates were then aggregated by calculating the mean. We obtained 18 datasets, one per condition, *i.e.* the combination of a data type (CP or L1000), a cell line (U2OS, A549 or MCF7) and a time point (6h, 24h, 48h), and containing 106 profiles corresponding to 106 compounds. Finally, we retrieved compound annotations gathering information about their modes of action and targets from the GitHub repository of the CP profiles available at https://github.com/broadinstitute/lincs-cell-painting^26^. The details of the 106 compounds are available in Table S3.

#### UMAP visualizations

Uniform manifold Approximations and Projections (UMAPs)^50^ were computed to visualize our datasets in a two-dimensional space, and study cell line and time effects on CP and L1000 data. UMAP is commonly used to reduce dimension because it is known to preserve both local and global structures of the data, meaning that the distances between distant points will be preserved as well as those between nearby points. We used the implementation of the umap-learn (v0.5.1) package of Python (v3.9.16^48^), with the default parameters (metric=euclidean, n_neighbors=15, min_dist=0.1).

#### Computation of cosine similarity between profiles

We calculated cosine similarity between the CP profiles of every condition on one side, and between the L1000 profiles of every condition on the other side. We then created heatmaps based on the cosine similarity matrix for both CP and L1000 using the seaborn^51^ package (v0.12.2) of Python.

#### Creation of modules of co-expressed genes and co-perturbed features

Weighted Gene-expression Correlation Network Analysis (WGCNA) is a method originally designed to identify groups of highly connected genes. The principle is based on correlation networks: a network is constructed based on gene expression profiles across samples, representing the interconnectivity between genes. The network is coded by an adjacency matrix, which can be likened to a correlation matrix between the genes. Clustering of this matrix allows for the identification of clusters of highly correlated genes, also known as modules^52^. We used the WGCNA package (v1.72-5) delivered by Langfelder et al.^52^ and implemented in R (v4.3.2^53^) to create modules of genes for each L1000 datasets. Modules are, by default, named by color. Since modules were created independently for each condition, two modules in different conditions can have the same color name, even if they are different.

WGCNA usually takes log fold changes or Z-scores as input for differential expression measures. As Cell Painting features are also represented as differential expression measures (Z-scores), we assumed that WGCNA can be extrapolated to Cell Painting to create modules of highly correlated features. For both CP and L1000, we chose signed networks, meaning that signs of correlation were considered. Then, in order to reduce noise in the adjacency matrix and to better approximate our network to a scale-free network (where some genes/features have more connections than others), we utilized the *pickSoftThreshold* function to select a soft power (between 1 and 30) for each dataset. Soft power refers to a parameter that controls the strength of the correlation between genes/features when constructing co-expression networks. It allows us to identify meaningful gene/feature modules by emphasizing strong correlations while downweighting weaker ones. Low soft powers can be associated with a high connectivity of genes/features in the network, resulting in very correlated and larger modules, which could invalidate the approximation with a scale-free network. We verified that the mean connectivity associated with the chosen soft power approached a few hundred at most. We set the minimum module size to 5 and merged modules that were correlated by more than 90%.

Module composition of each dataset is detailed in the Supplementary Table 1.

#### Customized enrichment between compounds and modules of genes or features

In order to link compounds from each dataset to the modules of genes and features they most significantly affected, we conducted custom enrichment analysis using the R EnrichIntersect package (v0.6)^54^ and calculated enrichment scores between each compound and module in each dataset. Scores were normalized. To evaluate the significance of the enrichment scores, we conducted 200 permutation tests, adjusted the p-values using the Bonferroni correction, and considered p-values < 0.05 to be significant.

We assumed that compounds perturbing the same modules of morphological features in a similar manner could also perturb the same modules of genes in a similar manner. Therefore, we clustered the compounds from each dataset individually based on the enrichment scores to group compounds that perturb the same modules. Subsequently, we used tanglegrams to compare the compound clustering obtained with both Cell Painting and L1000, in the 3 cell lines, at 6h and 48h. All analyses were conducted using the dendextend package in R (v1.17.1)^55^. The distance matrices between the enrichment scores were calculated using the euclidean distance, and the ward.D2 method was used for clustering. These analyses were not conducted for the 24h exposure time due to the inapplicability of WGCNA on the CP datasets of A549 and MCF7 under this time condition and resulting in the absence of modules.

#### Identification of modules of features and genes significantly representative of a core of HDAC or CDK inhibitors

We define as a core a group of compounds (HDAC or CDK inhibitors) that are found grouped together with a maximum distance of 6, whether clustering based on the enrichment scores of these compounds for the modules of features or the modules of genes, and in at least 2 out of 3 cell lines. The cores were defined using the tanglegrams of the 3 cell lines for the 6-hour time point. These same cores were then used to study their conservation at 48 hours. The core of HDAC inhibitors included vorinostat, panobinostat, resminostat, JNJ-26481585, belinostat and dacinostat. The core of CDK inhibitors comprised R547, PHA-793887, AZD5438, dinaciclib, AT-7519 and alvocidib.

To identify the modules of morphological features and genes that are significantly representative of either the HDAC inhibitors or the CDK inhibitors, we performed a Mann-Whitney test using the greater alternative for each cell line and duration of exposure. For each module, we compared the enrichment scores of the compounds in the respective core (HDAC or CDK) to those of the other compounds. Modules with a p-value < 0.05 were selected. We then evaluated the overlap of the selected modules of features and genes across cell lines and time conditions to create typical profiles of HDAC inhibitors and CDK inhibitors over time and cell lines. The overlaps were visualized using the upsetplot package in Python (v0.9.0)^56^.

#### Pathway enrichment analysis

We performed over-representation analysis to identify biological pathways and cellular components associated with different gene sets disturbed by the compounds of the core of HDAC or CDK inhibitors. We used the *enrichr* function of the GSEApy package (v1.1.0)^57^ and selected the Gene Ontology (GO) GO_Biological_Process_2021 and GO_Cellular_Component_2021 libraries and the human organism. GO terms with an adjusted p-value < 0.05 were considered significantly enriched.

