## Supplementary file for "Cell morphology and gene expression: tracking changes and complementarity across time and cell lines"

### Supplementary Figures

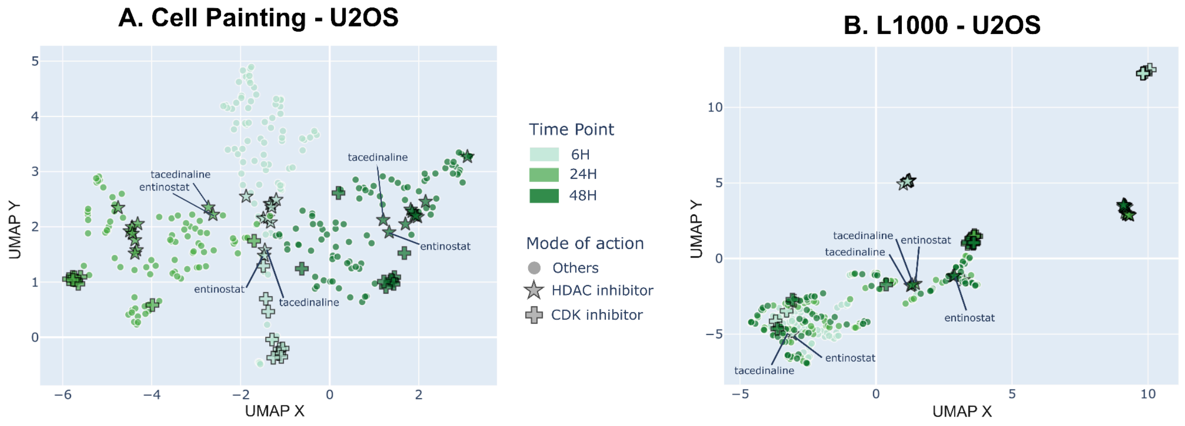


**Figure S1. Comparison over time of the phenotypic and transcriptomic signatures of HDAC and CDK inhibitors in the U2OS cell line.** Each point represents a compound tested in either CP **(A)** or L1000 **(B)** on U2OS cells. The colors transition from lighter to darker as the exposure time increases, going from 6h to 48h. HDAC inhibitors and CDK inhibitors are respectively symbolized by stars and crosses outlined in black, and the other compounds by circles. Points corresponding to entinostat and tacedinaline were specifically labeled.

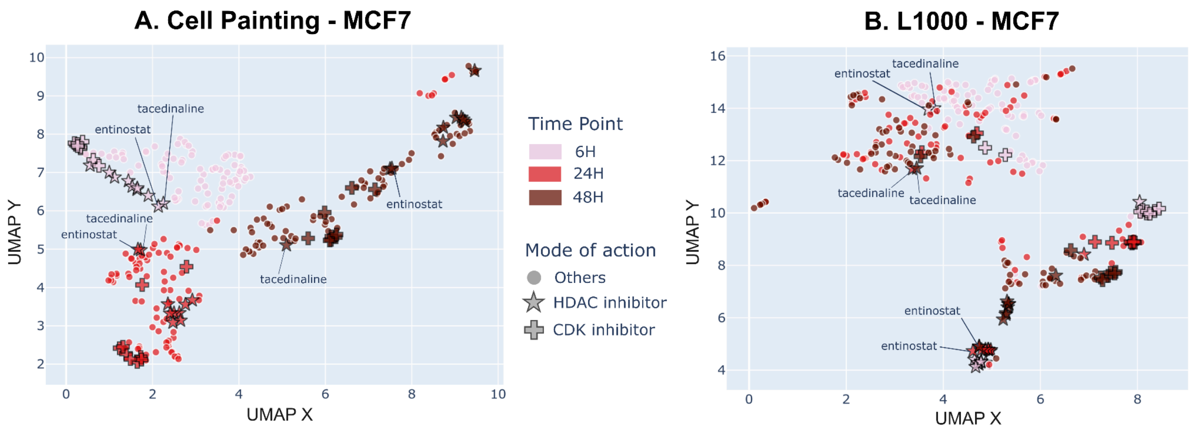


**Figure S2. Comparison over time of the phenotypic and transcriptomic signatures of HDAC and CDK inhibitors in the MCF7 cell line.** Each point represents a compound tested in either CP **(A)** or L1000 **(B)** on MCF7 cells. The colors transition from lighter to darker as the exposure time increases, going from 6h to 48h. HDAC inhibitors and CDK inhibitors are respectively symbolized by stars and crosses outlined in black, and the other compounds by circles. Points corresponding to entinostat and tacedinaline were specifically labeled.

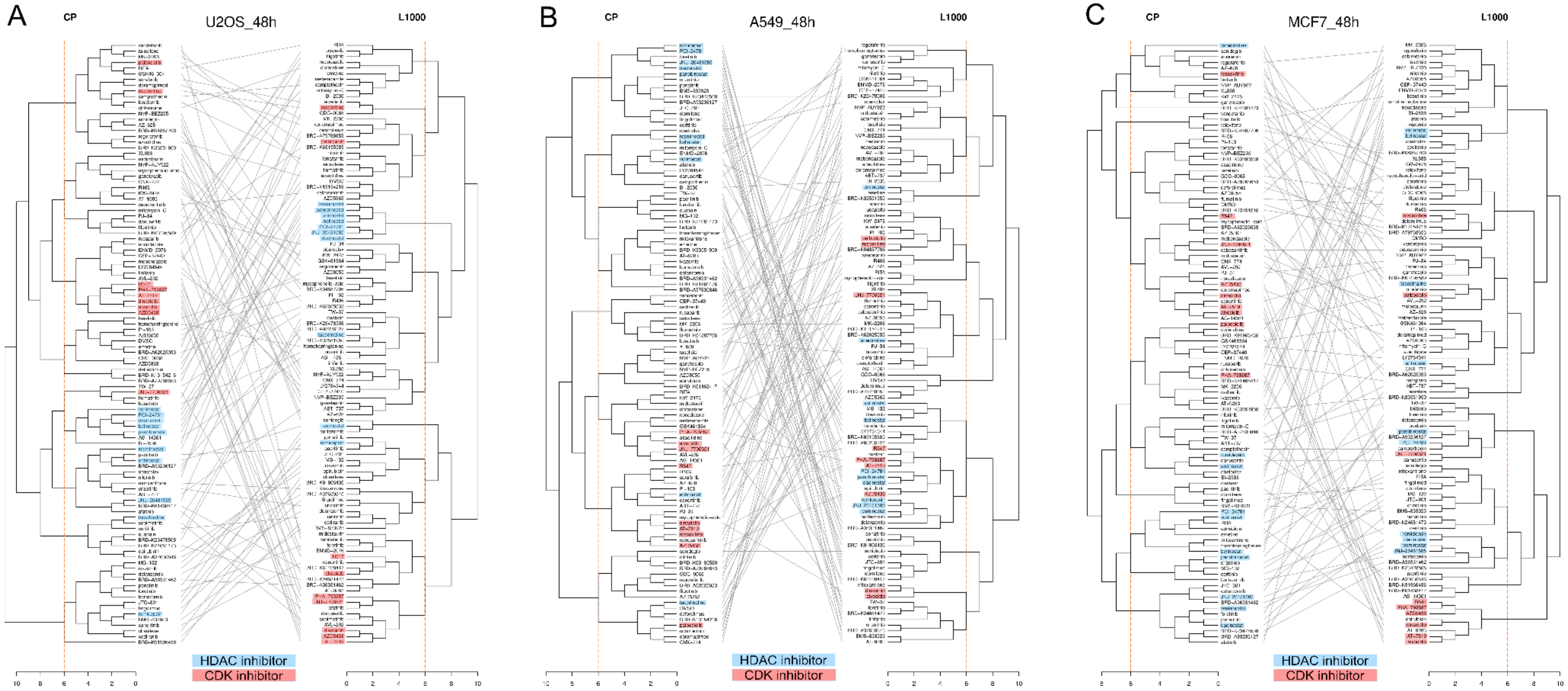
A
CP
U2OS_48h
L1000
HDAC inhibitor
CDK inhibitor
B
CP
A549_48h
L1000
HDAC inhibitor
CDK inhibitor
C
CP
MCF7_48h
L1000
HDAC inhibitor
CDK inhibitor

**Figure S3. Tanglegrams comparing the hierarchical clustering of compounds at the 48h time point in the U2OS, A549, and MCF7 cell lines.** Individual dendrograms were computed separately using distance matrices calculated with Euclidean distance and clustered with the ward.D2 method. A distance scale is located at the bottom of each dendrogram. The left dendrograms represent the clustering of compounds based on their customized enrichment scores for modules of CP features, while the right dendrograms represent clustering based on the same enrichment scores for modules of L1000 genes. Then, tanglegram visualization enables comparison between CP and L1000 dendrograms, where each leaf is a compound tested at 48h, for U2OS **(A)**, A549 **(B)** and MCF7 **(C)**. HDAC inhibitors are highlighted in blue, and CDK inhibitors in red. The vertical orange line at a clustering distance of 6 was used to define HDAC and CDK inhibitor cores as described in the Methods.
